# Short-term reciprocity in macaque’s social decision-making

**DOI:** 10.1101/695916

**Authors:** S. Ballesta, G. Reymond, J-R. Duhamel

**Author notes:** **Correspondence:** Dr. Sébastien Ballesta.

## Abstract

Primates live in complex social environments, where individuals create meaningful networks by adapting their behavior according to past experiences with others. Although free-ranging primates do show signs of reciprocity, experiments in more controlled environments have mainly failed to reproduce such social dynamics. Hence, the cognitive and neural processes allowing monkeys to reciprocate during social exchanges remains elusive. Here, pairs of long-tailed macaques (*Macaca fascicularis*) took turns into a social decision task involving the delivery of positive (juice reward) or negative (airpuff) outcomes. By analyzing the contingencies of one partner’s past decisions on the other’s future decisions, we demonstrate the presence of reciprocity, but only for the exchange of negative outcomes. Importantly, to display this decisional bias, the monkey needs to witness its partner’s decisions, since non-social deliveries of the same outcome did not have such effect. Withholding of negative outcomes also predicted future social decisions, which suggest that the observed tit-for-tat strategy may not only be motivated by retaliation after receiving an airpuff but also by the gratefulness of not having received one. These results clarify the apparent dichotomy within the scientific literature of reciprocity in non-human primates and suggest that their social cognition comprise revenge and gratitude.

## Introduction

During exchanges of positive or negative actions between individuals, prior outcomes may influence further social interactions. Many studies show that free-ranging primates display reciprocity in a range of social behaviors but reciprocal behavior has rarely been observed for food exchanges (Packer, 1977; Seyfarth and Cheney, 1984; de Waal and Luttrell, 1988; Ventura et al., 2006; Schino, 2007; Schino and Pellegrini, 2009; Carne et al., 2011; Weinstein and Capitanio, 2012; Xia et al., 2013; Amici et al., 2014; Borgeaud and Bshary, 2015; Molesti and Majolo, 2015, 2017). Surprisingly, under experimentally controlled conditions, monkeys failed to display signs of reciprocity (Brosnan et al., 2009; Claidière et al., 2015; Pelé et al., 2010; Suchak and de Waal, 2012; Yamamoto and Tanaka, 2009). This apparent lack of valuation of prior exchanges might be interpreted in several ways, such as real absence of short-term reciprocity, specific insensitivity to food exchanges, or poor understanding of their own and/or other’s agency inside such experimental apparatus (Drayton and Santos, 2014; Jaeggi et al., 2013). Hence, previous experimental studies may have underestimated the presence of reciprocity in non-human primates (Schweinfurth and Call, 2019). Here, we assessed the presence of reciprocity in well-controlled social decisions between young male long-tailed macaques (*Macaca fascicularis*). We designed a task where two monkeys faced each other, and alternately made social decisions involving juice or airpuff delivery to the partner or to an empty space (called ‘nobody’ hereafter) (Ballesta and Duhamel, 2015). In this task, the actor receives the same amount of juice drops after each of its social decisions, whether or not it involves positive (juice) or negative (airpuff) outcomes for its partner. Thus, choice in social decisions only differed as whether these would affect the partner or not, while the benefit for self was unchanged regardless of the choice. Importantly, each experimental session includes control non-social decisions where the monkey had to choose between the delivery of an outcome to himself or to nobody. It should be noted that the delivery of an airpuff is a negative somatosensory experience which can be compared to other negative stimuli used in experiments on social exchanges in humans (Shergill et al., 2003). The interpretation of its significance by non-human primates may be more straightforward than that of a food (or juice) exchange, which is, for instance, affected by several other factors such as satiety, spatial proximity of peers, and social hierarchy (Watson and Caldwell, 2009).

## Materials and Methods

### Animals

Four non-kin juvenile male long-tailed macaques (*Macaca fascicularis*) (aged 3+/-0.15 years, weight 5.7+/-0.8) were used as subjects. They were housed as a mini-colony in a relatively large enclosure (15m^3^) allowing direct physical interactions but also isolation through a system of sliding partitions. When isolated, monkeys could communicate visually and vocally. Animals were fed with monkey chow, fresh fruits and vegetables. The monkeys were maintained under scheduled access to fluid in order to maintain optimal motivation for juice. Extra fluid and fruits could be given as needed at the end of each day to maintain the animals’ proper fluid balance. In addition, animals had at least one day of free access to water each week. Animals were weighted before each experimental session. The difference in weight of the animals between periods of *ad libitum* and restricted access to fluids was always inferior to the ethical limit of 10% (which is in the range of what non-human primates can experience in the wild (Zurovsky et al., 1984)). According to recent scientific investigations, it is unlikely that such fluid restriction protocol had caused any physiological or psychological harm to the subjects (Gray et al., 2016). The cages were enriched with different toys, computer-based enrichment and substrates that promote social play, curiosity, object manipulation and foraging. This study was part of a larger project on social cognition involving neurophysiological and eye movement recordings which required specific surgical procedure to implant a head restraining device. Despite that this surgical procedure was not needed for this particular study, it does not impede the scientific validity of our results. In fact, in many other ethically accepted social neuroscience studies, non-human primates has already proven their ability to express coherent socio-cognitive behaviors (Chang et al., 2011; Haroush and Williams, 2015; Yoshida et al., 2011) that were consistent with naturally occurring social behaviors (Ballesta and Duhamel, 2015). Briefly, after a single sterile surgery performed under isoflurane anesthesia, the monkeys were then left to recover for at least one month with the proper antibiotic coverage, and pain-relievers were given as needed. All experimental procedures were approved by the animal care committee (Department of Veterinary Services, Health & Protection of Animals, permit number 69 029 0401) and the Biology Department of the University Claude Bernard Lyon 1, in conformity with the European Community standards for the care and use of laboratory animals [European Community Council Directive No. 86–609].

### Behavioral Procedures

The setup was designed to allow neurophysiological and eye movement recordings while the two monkeys interacted visually with each other and made behavioral choices using the touch panel interface (Fig. 1). Animal training and social decision task procedures are the same as in (Ballesta and Duhamel, 2015). Briefly, juice and airpuffs (4 bars) were delivered using a gravity-based solenoid device (Crist Instruments) and pressure gauge. Using a video projector and two semi-transparent mirrors (Beam splitter, 30% Reflection, 70% transmission, Edmund optics Inc.), the same visual stimuli was virtually projected in the visual plane of the two touch panels. Trials began with the appearance of a central colored square stimulus, which specified the actor monkey on that trial. When touched, this stimulus triggered the appearance of two distinct cues, each associated with a unique set of outcomes to the actor (self), the partner (other) or nobody (Fig. 1a). After 500ms, the square target was extinguished, and the monkey made its choice by touching the corresponding cue. After 1500ms, the outcomes were delivered. Different choice configurations were presented on successive trials from a pre-defined set of 4 possible offers, in randomly interleaved order. The two monkeys alternated as actor and partner on successive blocks of 30 seconds during which it performed an average of 3.14 (±0.9) trials. Unique sets of visual cues were associated with the outcomes delivered to each monkey. Behavioral control and visual displays were under the control of PCs running the REX/VEX system (Hays et al., 1982). All analog and digital data were logged and synchronized using Spike2 (Cambridge Electronic Design).

**Figure 1.**
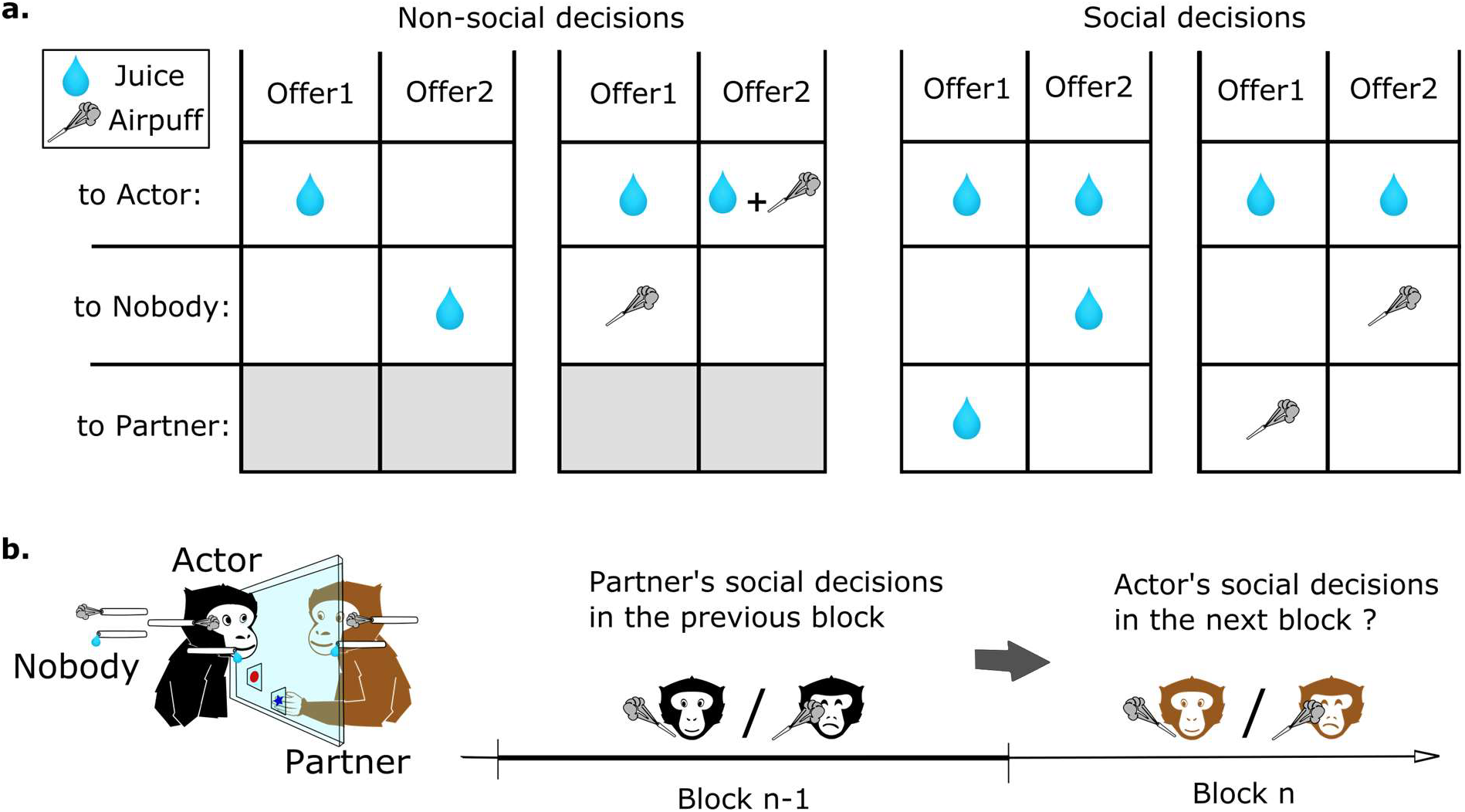
Experimental task and analysis to measure reciprocity during social exchanges. **(a)** Set of outcomes used for non-social and social decisions. Different offers configurations were presented on successive trials from a pre-defined set of 4 possible types of decisions, in randomly interleaved order. Each cue shape was associated with a unique set of outcomes to the actor (self), the partner (other) or nobody **(b)** Method used to analyze the effect of prior decisions on next decisions. The two monkeys alternated as actor and partner on successive blocks of 30 seconds. We compared the decisions of an actor monkey depending on the nature of the decisions of the other monkeys towards him on the preceding block of trials.

### Data analysis

Data from 49 experimental sessions (14,356 social and non-social decisions) where the actor and the partner monkey both performed more than 60 trials were analyzed using custom scripts written in Matlab R2015a (The Mathworks). We compared the decisions of an actor monkey depending on the nature of the decisions of the other monkeys towards him on a preceding block of trials (Fig. 1b). We computed a score for each preceding block by calculating the difference of each type of outcomes (for instance the difference in the number of airpuffs delivered to the partner or to nobody for each social decision of this kind). The computation of the previous block score (*pbs*) can be expressed by the following equation:

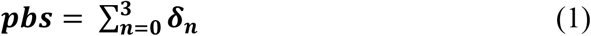

With δ being the outcome of a decision in each trial of the previous block and *n* the number of time that this type of decision was proposed in the previous block. Note that δ was equal to +1 and −1 when a prosocial or antisocial outcome was chosen, respectively (Fig. 2ab). To control for the non-social mere effect of receiving an airpuff, the same analysis was performed using the number of airpuff delivered to the actor by the actor himself in preceding block (Fig. 2c). A similar score was computed when considering the number of each outcome delivered in the preceding block (Fig. 2d). For a given type of decision, actor’s mean decision rates (*dr*) were calculated as follow for each previous block score:

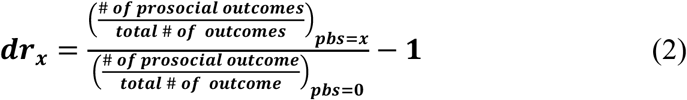

*x* is the value of the previous block score. As the decision type was drawn randomly on each trial, each block contained between 0 and 4 decisions of each type. Hence, *x* could vary between −4 and 4 but was constrained between −3 and 3 as extremes data points were not present in most of the experimental sessions. Note that a score equal to zero also includes blocks where this type of decision was absent from the previous block. A positive or negative value of *dr* indicate, respectively, an increase or decrease of the prosocial decisional tendencies of the actor. Linear fittings were performed in Matlab using the function *fitglme* and always considered actors’ identity as a random factor.

**Figure 2.**
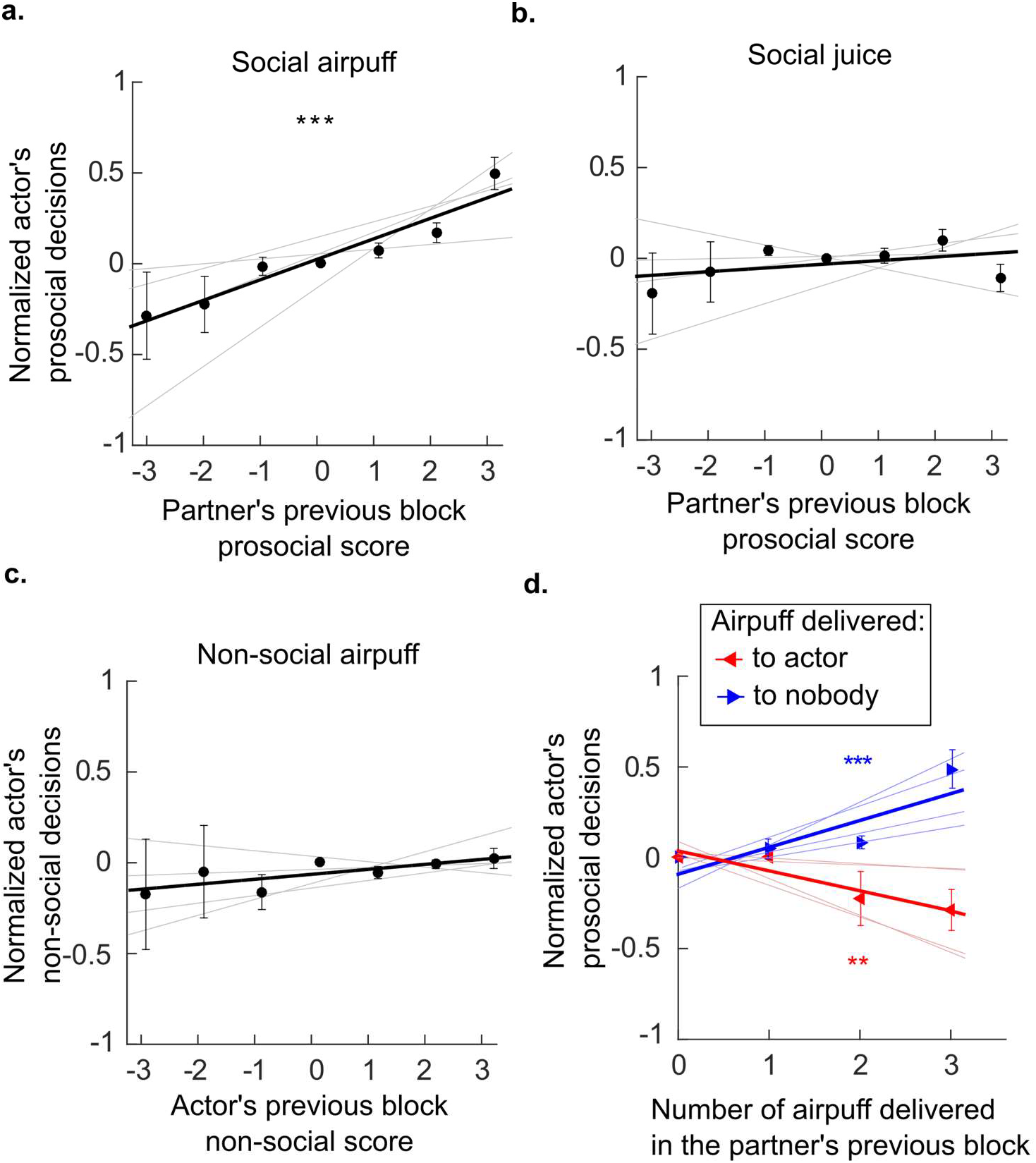
Previous delivery and avoidance of negative outcomes influenced subsequent social decisions. We assess whether prior decisions of a monkey (the partner or the actor) can influence the actor’s subsequent decisions. **(a)** Reciprocity between the social exchanges of airpuff (Generalized Linear Mixed Effect Model, R^2^=0.57, F=34.6, p<0.001). **(b)** Absence of reciprocity between the social exchanges of juice reward (GLME, R^2^=0.08, F=1.63, p=0.21). **(c)** Non-social delivery of airpuff does not influence social decision involving airpuff (GLME, R^2^=0.04, F=1.16, p=0.29). **(d)** Previous delivery and avoidance of airpuff predicted subsequent social decisions. (Number of airpuff delivered to nobody,GLME, R^2^=0.57, F=21.3, p<0.001; Number of airpuff delivered to the partner, GLME, R^2^=0.46, F=10.3, p=0.006). **(a-d)** Actors mean decision rates were normalized so that positive or negative value indicates, respectively, an increase or decrease of the prosocial decisional tendencies of the actor (see methods). Error bars represent SEM. Gray lines represent individual’s regressions.

## Results

Results were analyzed for all the possible pairs composed by 4 young male long-tailed macaques, who switched roles as actor and partner every 30 seconds during a social decision task (Fig. 1). The apparatus and the task were completely symmetric allowing considering the behavior of both of the animals during the 49 sessions, which represent 14,356 decisions. The decisions of the subjects were on average rational, for decision concerning self only, and prosocial, for decisions implicating the partner. The monkeys preferred to grant and avoid airpuffs (97% ±2, 90% ±3, Wilcoxon signed rank test p<0.01) to self, and showed a similar, though less pronounced, tendency for the same outcomes to their partners (68% ±3 and 60% ±4 for juice and airpuff respectively, Wilcoxon signed rank test p<0.01). In order to assess the presence of contingency within decisions of each subject, we measured the effect of its own decisions in the prior block (i.e. made within at most 60 seconds) onto its next decisions. In addition, to assess the presence of contingency between decisions of the two subjects, we measured the effect of the other monkey’s decisions in the prior block on the actor next decisions. Each previous block of partner’s social decisions was scored by computing the difference in the number of each outcome for a given type of decision (see methods). Generalized linear mixed effect models (GLME, see methods) were computed to test whether the partner’s previous decisions was a significant predictor of the actor’s subsequent decisions (Fig. 2). This analysis revealed that the partner’s social decision involving airpuff delivery in the previous block was a significant predictor of the actor’s social decisions of the same type in the current block (Fig. 2a, GLME, R^2^=0.57, F=34.6 p<0.001). However such relation was not found when the social decisions involved juice reward (Fig. 2b, GLME, R^2^=0.04, F=1.1, p=0.31) or when the social decisions of the partner’s and actors were of different type (past social decision: airpuff, current social decision: reward, GLME, R^2^=0.08, F=1.63, p=0.21; past social decision: reward, current social decision: airpuff, GLME, R^2^=0.12, F=3.8, p=0.07). Importantly, to control for the mere effect of receiving an airpuff independently from the identity of the sender, we also compared the next social decisions of the actor according to its own prior non-social decisions (e.g. whether the actor had sent airpuff to its own face on a prior block), and did not find significant results (Fig. 2c, GLME, R^2^=0.04, F=1.16, p=0.29). This result suggests that having recently received an airpuff (which could plausibly make the monkey irritable, irrational, or less attentive to the partner’s fate) is not by itself sufficient to increase the delivery of this aversive outcome to the partner. Finally, we also scored the previous blocks using only the number of a given outcomes delivery and performed similar analysis. We found that both the numbers of airpuffs delivered to nobody and to the actor by the partner were significant predictors of the actor’s social decisions involving airpuff deliveries (Fig. 2d, Number of airpuffs delivered to nobody: GLME, R^2^=0.57, F=21.3, p<0.001; Number of airpuffs delivered to the partner: GLME, R^2^=0.46, F=10.3, p=0.006). In other words, the number of airpuffs received and also avoided by the actor due to its partner decisions in the previous block influenced his own social decision in the next block. Similar analysis using the number of the outcomes from other types of decision was not significantly predicting the actors social decisions (Number of Juice delivered to the partner: GLME, R^2^=0, F=0.09, p=0.76; number of juice delivered to nobody: GLME, R^2^=0.15, F=1, p=0.32; number of airpuff delivered to the actor by the actor: GLME, R^2^=0.13, F=2.47, p=0.14; number of airpuff delivered to nobody by the actor: GLME, R^2^=0, F=0, p=0.94).

## Discussion

Our results show that during dynamic exchanges, macaques’ social decisions can be influenced by the prior social decisions of a peer. In order to control for a potential non-social confound, we have assessed the influence of a prior self-induced airpuff on a future social decision and found no effect. This ruled out trivial explanations of our results implying that receiving an airpuff increased the monkey’s arousal, which in turn would have negatively affected its prosocial motivation or its general ability to make decisions. This suggests that witnessing the other monkey’s social decisions is necessary to induce the observed social decision modulations. Other alternative interpretations of our results can be considered. For instance, one could argue that the observed reciprocity was due to a mirroring of the partner’s previous actions (Suchak and de Waal, 2012). However, the actions themselves (i.e. pointing gesture toward an icon on a touch screen) were the same for all choices and the visual cues associated with each outcome were specific for each monkey and each type of decisions. Hence, advocating for such form of reciprocity would imply that the animals were not imitating the mere gesture of the partner (Nagasaka et al., 2013) or the choice of a specific visual stimulus but the action’s intended effect. Such line of interpretation would somehow relate our results to a social form of cognitive imitation (Subiaul et al., 2004). However, the absence of reciprocity for the delivery of juice reward rules out the idea that our results can be solely explained by a purely reflective mechanism. Despite our relatively modest sample size, our data analysis strongly support the idea that the observed decisional bias is explained by a non-reflexive use of others’ behaviors to guide choices, as observed by other neuroscience studies in laboratory settings (Haroush and Williams, 2015; Yoshida et al., 2011). Ethological reports have shown that macaques can reciprocate positive behavior, such as social grooming (Schino, 2007) and can exhibit revenge-like behavior (Aureli et al., 1992; Silk, 1992). Our results could be thus interpreted as a proof of macaques following a tit-for-tat strategy during experimental social exchanges. It should be noted that, on average, the monkeys were prosocial as they significantly preferred to refrain from causing harm to their partners (for more details see (Ballesta and Duhamel, 2015)). Therefore, tit-for-tat could not be a dominant strategy in the present case, as it should have led to an escalation of retaliatory actions and, in the long run, to a tendency toward antisocial decision-making. The use of retaliation was thus likely balanced by other cognitive processes such as the desire to preserve other’s welfare or social bonds (Ballesta and Duhamel, 2015; de Waal and Suchak, 2010; Yamamoto and Takimoto, 2012). Monkeys therefore might use retaliation parsimoniously which is consistent with comparable studies involving human subjects (Fitz et al., 1979, 1983). Additionally, tit-for-tat strategy could be interpreted differently as meaning that the monkey recognized its partner’s active airpuff withholding and thus increased it prosocial tendency as proof of gratefulness. At first, it seems easier to recognize the social motivation of others on the basis of the actual consequences of their action toward self (here, the delivery of an airpuff to the other monkey), compare to the absence of consequences (the delivery of an airpuff in an empty space). However, our results show that the number of airpuff delivery and avoidance are both sufficient to measure contingencies between social exchanges. This suggest that the observed tit-for-tat strategy may not only be motivated by retaliation after receiving an airpuff but also by the gratefulness of not having received one. This unexpected display of gratitude in a despotic non-human primate species calls for further investigations as this higher-order cognitive ability seems crucial to establish prosociality in humans (Ma et al., 2017). In fact, retaliation and gratitude are likely to depend on evolutionary rooted social cognition, as preverbal infants can identify the nature of the social motivation of individuals during the observation of social interactions (Hamlin et al., 2007; Kuhlmeier et al., 2003).

Monkeys can perform social exchange in different currencies (e.g. a female grooming a mother to gain access to her infant) (Ventura et al., 2006; Carne et al., 2011; Borgeaud and Bshary, 2015). In our study, we did not find any evidence of such abilities to transform a social exchange into another form of social exchange. This negative result could be simply explained by the fact that our macaques did not reciprocate juice reward delivery. Recent ethological report show the absence of short term contingencies between grooming and food tolerance in a different species of macaques (Molesti and Majolo, 2015) and in bonobos (Goldstone et al., 2016). This is somehow consistent with our results and underline the singularity of food sharing behaviors (Watson and Caldwell, 2009). It is thus legitimate to challenge the relevancy of food allocation tasks to study primates’ social behaviors (Carter, 2014; Jaeggi et al., 2013; Watson and Caldwell, 2009). It is unclear whether macaques can conceive the agency in a transfer of ownership, and whether this represents a relevant social cue. In macaques, the tolerance for co-feeding is unlikely to be bi-directional as it mainly occurs between the mother and her infant, or between a potent and a more subordinate individual (Dubuc et al., 2012; Jaeggi and Van Schaik, 2011; Massen et al., 2010; Ventura et al., 2006). In macaque despotic societies dominant individuals naturally have a “right of preemption” on the subordinates’ goods which likely made reciprocal exchanges of food irrelevant. Future studies should take these observations into consideration, especially when performances in juice reward allocation tasks are taken as a proxy of macaque’s social motivation in neuroscience (Azzi et al., 2012; Chang et al., 2011). To overcome these methodological and conceptual challenges and study exchanges of positive acts under experimental condition in primates, the use of grooming by the experimenters (or by a dedicated device) might be considered in further investigations (Grandi and Ishida, 2015; Taira and Rolls, 1996). To conclude, by using both positive and negative outcomes, this study has clarified the apparent dichotomy within the scientific literature of reciprocity in non-human primates. We found contingencies in the social exchange of negative acts in macaques and show that macaques can use tit-for-tat strategies and gratitude in their social exchanges which represent a new insight into the nature of non-human primates’ social cognition. These results lay the foundation for the investigations of the cognitive and neural basis of reciprocity in exchange of negative acts in non-human primates.

## Authors’ contributions

S.B., G.R. and J-R.D. performed analyses, discussed the results and contributed to the text of the manuscript. S.B performed the experiment. All authors reviewed the manuscript.

## Acknowledgements

We thank Mathieu Pozzobon for assistance in animal training, Serge Pinède for technical support. We also thank Jean-Luc Charieau and Fabrice Hérant for expert animal care.

## Datasets are available on request

The raw data supporting the conclusions of this manuscript will be made available by the authors, without undue reservation, to any qualified researcher.

## References

Amici, F., Aureli, F., Mundry, R., Amaro, A. S., Barroso, A. M., Ferretti, J., et al. (2014). Calculated reciprocity? A comparative test with six primate species. Primates J. Primatol. 55, 447–457. doi: 10.1007/s10329-014-0424-4.

Aureli, F., Cozzolino, R., Cordischi, C., and Scucchi, S. (1992). Kin-oriented redirection among Japanese macaques: an expression of a revenge system? Anim. Behav. 44, 283–291. doi: 10.1016/0003-3472(92)90034-7.

Azzi, J. C. B., Sirigu, A., and Duhamel, J.-R. (2012). Modulation of value representation by social context in the primate orbitofrontal cortex. Proc. Natl. Acad. Sci. U. S. A. 109, 2126–2131. doi: 10.1073/pnas.1111715109.

Ballesta, S., and Duhamel, J.-R. (2015). Rudimentary empathy in macaques’ social decision-making. Proc. Natl. Acad. Sci., 201504454.

Borgeaud, C., and Bshary, R. (2015). Wild Vervet Monkeys Trade Tolerance and Specific Coalitionary Support for Grooming in Experimentally Induced Conflicts. Curr. Biol. 25, 3011–3016. doi: 10.1016/j.cub.2015.10.016.

Brosnan, S. F., Silk, J. B., Henrich, J., Mareno, M. C., Lambeth, S. P., and Schapiro, S. J. (2009). Chimpanzees (Pan troglodytes) do not develop contingent reciprocity in an experimental task. Anim. Cogn. 12, 587–597. doi: 10.1007/s10071-009-0218-z.

Carne, C., Wiper, S., and Semple, S. (2011). Reciprocation and interchange of grooming, agonistic support, feeding tolerance, and aggression in semi-free-ranging Barbary macaques. Am. J. Primatol. 73, 1127–1133. doi: 10.1002/ajp.20979.

Carter, G. (2014). The Reciprocity Controversy. Anim. Behav. Cogn. 1, 368. doi: 10.12966/abc.08.11.2014.

Chang, S. W. C., Winecoff, A. A., and Platt, M. L. (2011). Vicarious Reinforcement in Rhesus Macaques (Macaca Mulatta). Front. Neurosci. 5.

Claidière, N., Whiten, A., Mareno, M. C., Messer, E. J. E., Brosnan, S. F., Hopper, L. M., et al. (2015). Selective and contagious prosocial resource donation in capuchin monkeys, chimpanzees and humans. Sci. Rep. 5, 7631. doi: 10.1038/srep07631.

de Waal, F. B. M., and Luttrell, L. M. (1988). Mechanisms of social reciprocity in three primate species: Symmetrical relationship characteristics or cognition? Ethol. Sociobiol. 9, 101–118. doi: 10.1016/0162-3095(88)90016-7.

de Waal, F. B. M., and Suchak, M. (2010). Prosocial primates: selfish and unselfish motivations. Philos Trans R Soc Lond B Biol Sci 365, 2711–2722. doi: http://dx.doi.org/10.1098/rstb.2010.0119.

Drayton, L., and Santos, L. (2014). Insights into Intraspecies Variation in Primate Prosocial Behavior: Capuchins (Cebus apella) Fail to Show Prosociality on a Touchscreen Task. Behav. Sci. 4, 87–101. doi: 10.3390/bs4020087.

Dubuc, C., Hughes, K. D., Cascio, J., and Santos, L. R. (2012). Social tolerance in a despotic primate: co-feeding between consortship partners in rhesus macaques. Am. J. Phys. Anthropol. 148, 73–80. doi: 10.1002/ajpa.22043.

Fitz, D., Kimble, C., and Heidenfelder, K. (1979). Effects of external incentive and aggressive predisposition on aggression reduction. J. Psychol. 103, 71–80. doi: 10.1080/00223980.1979.9915116.

Fitz, D., Marwit, S. J., and Gerstenzang, S. (1983). Hostility Reduction In Married and Unacquainted Couples. J. Psychol. 115, 177–184. doi: 10.1080/00223980.1983.9915433.

Goldstone, L. G., Sommer, V., Nurmi, N., Stephens, C., and Fruth, B. (2016). Food begging and sharing in wild bonobos (Pan paniscus): assessing relationship quality? Primates J. Primatol. 57, 367–376. doi: 10.1007/s10329-016-0522-6.

Grandi, L. C., and Ishida, H. (2015). The Physiological Effect of Human Grooming on the Heart Rate and the Heart Rate Variability of Laboratory Non-Human Primates: A Pilot Study in Male Rhesus Monkeys. Front. Vet. Sci. 2. doi: 10.3389/fvets.2015.00050.

Gray, H., Bertrand, H., Mindus, C., Flecknell, P., Rowe, C., and Thiele, A. (2016). Physiological, Behavioral, and Scientific Impact of Different Fluid Control Protocols in the Rhesus Macaque (Macaca mulatta). eNeuro 3, ENEURO.0195-16.2016. doi: 10.1523/ENEURO.0195-16.2016.

Hamlin, J. K., Wynn, K., and Bloom, P. (2007). Social evaluation by preverbal infants. Nature 450, 557–559. doi: 10.1038/nature06288.

Haroush, K., and Williams, Z. M. (2015). Neuronal prediction of opponent’s behavior during cooperative social interchange in primates. Cell 160, 1233–1245. doi: 10.1016/j.cell.2015.01.045.

Hays, A., Richmond, B., and Optican, L. (1982). Unix-based multiple-process system, for real-time data acquisition and control. Electron Conventions,El Segundo, CA.

Jaeggi, A. V., De Groot, E., Stevens, J. M. G., and Van Schaik, C. P. (2013). Mechanisms of reciprocity in primates: testing for short-term contingency of grooming and food sharing in bonobos and chimpanzees. Evol. Hum. Behav. 34, 69–77. doi: 10.1016/j.evolhumbehav.2012.09.005.

Jaeggi, A. V., and Van Schaik, C. P. (2011). The evolution of food sharing in primates. Behav. Ecol. Sociobiol. 65, 2125–2140. doi: 10.1007/s00265-011-1221-3.

Kuhlmeier, V., Wynn, K., and Bloom, P. (2003). Attribution of dispositional states by 12-month-olds. Psychol. Sci. 14, 402–408.

Ma, L. K., Tunney, R. J., and Ferguson, E. (2017). Does gratitude enhance prosociality?: A meta-analytic review. Psychol. Bull. 143, 601–635. doi: 10.1037/bul0000103.

Massen, J. J. M., van den Berg, L. M., Spruijt, B. M., and Sterck, E. H. M. (2010). Generous leaders and selfish underdogs: pro-sociality in despotic macaques. PloS One 5, e9734. doi: 10.1371/journal.pone.0009734.

Molesti, S., and Majolo, B. (2015). No Short-Term Contingency Between Grooming and Food Tolerance in Barbary Macaques (Macaca sylvanus). Ethology 121, 372–382. doi: 10.1111/eth.12346.

Molesti, S., and Majolo, B. (2017). Evidence of direct reciprocity, but not of indirect and generalized reciprocity, in the grooming exchanges of wild Barbary macaques (Macaca sylvanus). Am. J. Primatol. 79, e22679. doi: 10.1002/ajp.22679.

Nagasaka, Y., Chao, Z. C., Hasegawa, N., Notoya, T., and Fujii, N. (2013). Spontaneous synchronization of arm motion between Japanese macaques. Sci. Rep. 3. doi: 10.1038/srep01151.

Packer, C. (1977). Reciprocal altruism in Papio anubis. Nature 265, 441–443. doi:10.1038/265441a0.

Pelé, M., Thierry, B., Call, J., and Dufour, V. (2010). Monkeys fail to reciprocate in an exchange task. Anim. Cogn. 13, 745–751. doi: 10.1007/s10071-010-0325-x.

Schino, G. (2007). Grooming and agonistic support: a meta-analysis of primate reciprocal altruism. Behav. Ecol. 18, 115–120.

Schino, G., and Pellegrini, B. (2009). Grooming in mandrills and the time frame of reciprocal partner choice. Am. J. Primatol. 71, 884–888. doi: 10.1002/ajp.20719.

Schweinfurth, M. K., and Call, J. (2019). Revisiting the possibility of reciprocal help in non-human primates. Neurosci. Biobehav. Rev. doi: 10.1016/j.neubiorev.2019.06.026.

Seyfarth, R. M., and Cheney, D. L. (1984). Grooming, alliances and reciprocal altruism in vervet monkeys. Nature 308, 541–543.

Shergill, S. S., Bays, P. M., Frith, C. D., and Wolpert, D. M. (2003). Two eyes for an eye: the neuroscience of force escalation. Science 301, 187–187.

Silk, J. B. (1992). The patterning of intervention among male bonnet macaques: reciprocity, revenge, and loyalty. Curr. Anthropol., 318–325.

Subiaul, F., Cantlon, J. F., Holloway, R. L., and Terrace, H. S. (2004). Cognitive imitation in rhesus macaques. Science 305, 407–410. doi: 10.1126/science.1099136.

Suchak, M., and de Waal, F. B. M. (2012). Monkeys benefit from reciprocity without the cognitive burden. Proc. Natl. Acad. Sci. 109, 15191–15196. doi: 10.1073/pnas.1213173109.

Taira, K., and Rolls, E. T. (1996). Receiving grooming as a reinforcer for the monkey. Physiol. Behav. 59, 1189–1192.

Ventura, R., Majolo, B., Koyama, N. F., Hardie, S., and Schino, G. (2006). Reciprocation and interchange in wild Japanese macaques: grooming, cofeeding, and agonistic support. Am. J. Primatol. 68, 1138–1149. doi: 10.1002/ajp.20314.

Watson, C. F. I., and Caldwell, C. A. (2009). Understanding Behavioral Traditions in Primates: Are Current Experimental Approaches Too Focused on Food? Int. J. Primatol. 30, 143–167. doi: 10.1007/s10764-009-9334-5.

Weinstein, T. A. R., and Capitanio, J. P. (2012). Longitudinal stability of friendships in rhesus monkeys (Macaca mulatta): Individual- and relationship-level effects. J. Comp. Psychol. 126, 97–108. doi: 10.1037/a0025607.

Xia, D.-P., Li, J.-H., Garber, P. A., Matheson, M. D., Sun, B.-H., and Zhu, Y. (2013). Grooming reciprocity in male Tibetan macaques. Am. J. Primatol. 75, 1009–1020. doi: 10.1002/ajp.22165.

Yamamoto, S., and Takimoto, A. (2012). Empathy and Fairness: Psychological Mechanisms for Eliciting and Maintaining Prosociality and Cooperation in Primates. Soc. Justice Res. 25, 233–255. doi: 10.1007/s11211-012-0160-0.

Yamamoto, S., and Tanaka, M. (2009). Do chimpanzees (Pan troglodytes) spontaneously take turns in a reciprocal cooperation task? J. Comp. Psychol. Wash. DC 1983 123, 242–249. doi: 10.1037/a0015838.

Yoshida, K., Saito, N., Iriki, A., and Isoda, M. (2011). Representation of Others’ Action by Neurons in Monkey Medial Frontal Cortex. Curr. Biol.

Zurovsky, Y., Shkolnik, A., and Ovadia, M. (1984). Conservation of blood plasma fluids in hamadryas baboons after thermal dehydration. J. Appl. Physiol. 57, 768–771.

